# Rule-based modeling using wildcards

**DOI:** 10.1101/112052

**Authors:** Steven S. Andrews

**Affiliations:** Fred Hutchinson Cancer Research Center, Seattle, WA, USA; Isaac Newton Institute for Mathematical Sciences, Cambridge, UK

**Keywords:** rule-based modeling, particle-based simulation, wildcards, reaction networks, spatial simulation, stochastic simulation, Brownian dynamics

## Abstract

Many biological molecules exist in multiple variants, such as proteins with different post-translational modifications, DNAs with different sequences, and phospholipids with different chain lengths. Representing these variants as distinct species, as most biochemical simulators do, leads to the problem that the number of species, and chemical reactions that interconvert them, typically increase combinatorially with the number of ways that the molecules can vary. This can be alleviated by “rule-based modeling methods,” in which software generates the chemical reaction network from relatively simple “rules.” This article presents a new approach to rule-based modeling. It is based on wildcards that match to species names, much as wildcards can match to file names in computer operating systems. It is much simpler to use than the formal rule-based modeling approaches developed previously but can also lead to unintended consequences if not used carefully. This article demonstrates rule-based modeling with wildcards through examples for: signaling systems, protein complexation, polymerization, nucleic acid sequence copying and mutation, the “SMILES” chemical notation, and others. The method is implemented in Smoldyn, a spatial and stochastic biochemical simulator, for both the generate-first and on-the-fly expansion, meaning whether the reaction network is generated before or during the simulation.

## 1 Introduction

Since about the time that Boyle posited that matter was composed of minute particles “associated into minute masses or clusters” (***1***), now recognized as molecules, the dominant paradigm in chemistry has been to classify molecules into chemical species. This paradigm forms the foundation of chemical kinetics (***2, 3***) and is supported by the finding that different molecules of the same species are completely indistinguishable from each other (***4, 5***). Correspondingly, most modern biochemical simulation software represents molecules as members of species, treating all members of a single species identically (see reviews (***6, 7***)). However, many biological molecules do not fit neatly into these classes. For example, a cell might have a hundred or more DNA molecules, each with a different sequence. Similarly, a cell might have thousands of copies of some protein, but the copies vary according to whether they are bound to other proteins, bound to cofactors, or post-translationally modified with phosphate, methyl, or other moieties.

Several approaches have been developed to represent this molecular variation in computational models. One is to represent every multimer as an explicit graph, including its component monomers and their interconnections (e.g. (***8-12***). Here, every molecule is its own entity and the concept of a species as a class of molecules is unnecessary. A second approach is to maintain the species concept, but to include states in the molecule definitions. For example, some biochemical simulators allow molecules to have modification states (***13-15***), surface-binding states (***16***), or an entire hierarchy of states (***17***). A third approach is to define each molecular variant as a separate species, with minimal variation within species. The possible variations can lead to a combinatorial expansion in the number of species (***18***), leading to the development of so-called *rule-based modeling* methods for automating reaction network expansion from “rules” that describe molecular complexation and modifications (e.g. (***19-21***)).

I recently followed this last approach, adding rule-based modeling to the Smoldyn simulator (***22***). Smoldyn is a widely used biochemical simulator that represents molecules as individual particles in 1D, 2D, or 3D space; these molecules diffuse, react with each other, and interact with surfaces (***16, 23, 24***). Smoldyn now supports two types of rule-based modeling. First, it sends any rules in the user’s input file that are written in the BioNetGen language (BNGL) to the BioNetGen software (***19, 25***). BioNetGen expands the rules to lists of species and reactions; then, Smoldyn reads the species and reactions, computes diffusion coefficients, graphical display parameters, and surface interactions for the new species, and runs the simulation (***22***). Second, Smoldyn performs a separate type of rule-based modeling using wildcard characters (***22***), which is the focus of this article. In this method, a modeler can specify groups of species using wildcard characters, much as a computer user can specify groups of files using wildcard characters. When used in chemical reactions, these wildcards can be used to define new species and new reactions.

Whereas rule-based modeling with formal languages, such as BNGL, were designed around an underlying model of how protein complexation and modification generally works, this is not the case for wildcards. Instead, the wildcard approach is simply a well-defined set of text-replacement tools with which the modeler can create his or her own notational scheme. This offers substantial versatility and generally simplifies input files. However, it can also lead to undesired results if not used carefully. Thus, the main text of this article focuses on the precise behavior of the wildcards, while the notes section presents examples that illustrate how the method can be used effectively.

## 2 Materials

Download Smoldyn from http://www.smoldyn.org. Smoldyn is free, open source, and licensed under the relatively permissive LGPL. The download package comes with install scripts, a detailed user’s manual, about 100 example input files, related software tools (including BioNetGen), and, if desired, the source code. Install on Macs and Windows with the install scripts, which is generally easy. Install on Linux computers by compiling the source code with CMake and Make, which is also straightforward. Smoldyn runs on most laptops and larger computers that are less than 5 years old, as well as many older computers. Support is available by e-mailing.

## 3 Methods

### 3.1 Running Smoldyn

To simulate a model in Smoldyn, start by describing the model in the Smoldyn language using a plain text file. Ref. (***26***) and the Smoldyn User’s Manual (included in the download package) describe how to write input files and give suggestions for parameter values.

Run Smoldyn at a shell prompt (a “Terminal” or “Command Line” application) by typing `smoldyn myfile.txt`, where `myfile.txt` is your configuration file name. Upon starting, Smoldyn reads model parameters from your configuration file, calculates and displays simulation parameters, and runs the simulation. As the simulation runs, Smoldyn displays the simulated system to a graphics window and saves quantitative data to one or more output files.

### 3.2 Wildcards for matching

Molecules in Smoldyn are classified into chemical species and can also adopt any of 5 physical *states.* These states are in solution (e.g. a cell’s cytoplasm) or the 4 surface-bound states called “front,” “back,” “up,” and “down.” Originally, the former two surface-bound states were for peripheral membrane proteins and the latter two were for integral membrane proteins, although they are all essentially equivalent in practice. All molecules of a single species and state behave identically, meaning that they have the same diffusion coefficients, graphical display parameters, surface interaction rates, and chemical reaction rates. Any other molecular variation needs to be expressed using separate species. For example, if a model includes the yeast Fus3 protein, which can bind to zero, one, or two phosphate groups (***27***), then each of its phosphorylation states would need to be represented as a separate species. Alternatively, if a model includes a receptor that diffuses at one rate in normal membrane regions and more slowly in lipid rafts, then this variation would again need to be represented using separate species.

Modeling such variations can lead to a rapid proliferation of separate species and so can become tedious to address. Use of wildcards alleviates this problem because it enables one to represent multiple species at once. For example, if the three Fus3 species were named Fus3, Fus3p, and Fus3pp, then the *species pattern* Fus3* would represent all three species. Also, if the receptors mentioned above were named R_normal and R_raft, then the species pattern R_* would represent both species. More generally, a species pattern is a species name that may include wildcard characters. In both of these examples, the ‘*’ wildcard is used to represent variable portions of the species names.

Smoldyn supports *text-matching* and *structural* wildcards, where the former ones match to specific portions of the species names and the latter ones enable logical operations in species patterns. The text-matching wildcards include ‘*’, which matches to any zero or more characters, ‘?’, which matches to any one character, and […], which matches to any one character from a specified list. The structural wildcard characters include ‘I’, which is an OR operator, ‘&’, which is a permutation operator, and {…}, which specifies the order of operation for the other two structural wildcards (the normal order of operations is that ‘&’ takes precedence over ‘I’). The structural wildcards are most easily explained through examples, this time using the generic protein monomer names A, B, and C: the pattern AIB matches to either A or B, the pattern A&B matches to either AB or BA, the pattern A&B|C matches to AB, BA, or C, and the pattern A&{B|C} matches to AB, BA, AC, or CA. See Table 1.

**Table 1.**
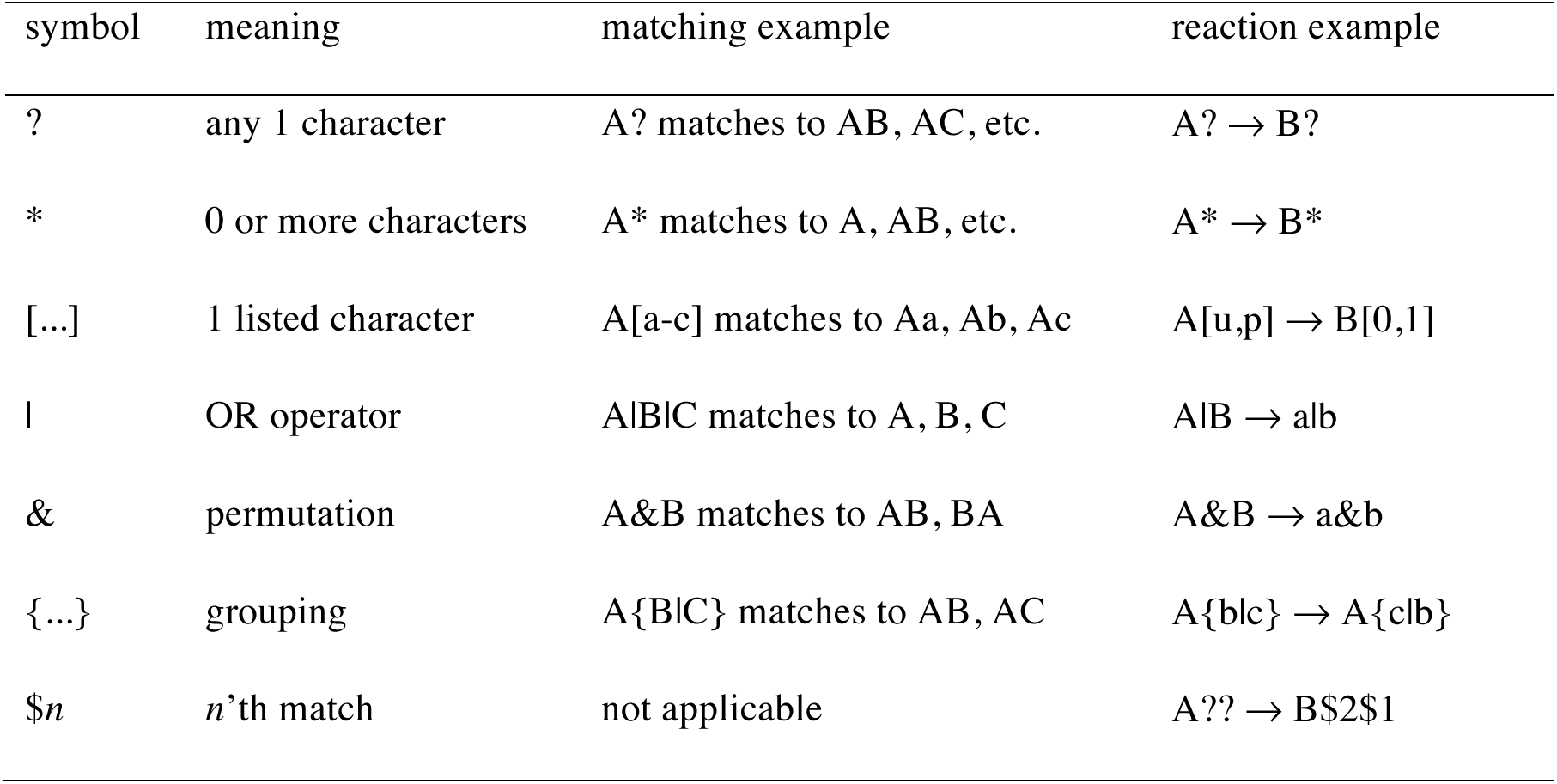
Smoldyn wildcards

Internally, when Smoldyn parses the user’s input file and expects a species name, it inputs the given text as a species pattern. The pattern may be as simple as a single species name but could also include one or more wildcard characters. If the pattern does not include structural wildcards, then it is an *elementary pattern.* On the other hand, if it does include structural wildcards, such as the pattern A&*, then Smoldyn first expands it to a list of *elementary patterns*; here, Smoldyn would expand the example to A* and *A. Next, Smoldyn scans through its list of species names to see which ones can match the elementary pattern(s). These matching species form a *species group.* If the pattern arose in a statement that defines species attributes (e.g. difc, for specifying the diffusion coefficient), then Smoldyn assigns the same attribute value to all species within the species group. Alternatively, if the pattern arose in a command that outputs information about molecules (e.g. molcountspecies, which counts the number of molecules of a given species or species group), then Smoldyn combines the appropriate information for all of the molecules that are in the species group.

### 3.3 Wildcards for substitutions

Smoldyn also supports wildcards in chemical reaction definitions, where they can be used to specify multiple chemical reactions at once. Smoldyn inputs each chemical reaction equations as a *reaction pattern,* which again may include wildcards but does not have to.

First, consider elementary reaction patterns, meaning reaction patterns that do not contain structural wildcards. In this case, Smoldyn substitutes any text that the wildcards match for the reactants into the corresponding wildcards in the products. For example, the reaction Ste5 + Fus3* → Ste5-Fus3* specifies that any of the three Fus3 species described above can associate with the Ste5 protein (***27***). In this case, the respective products would be Ste5-Fus3, Ste5-Fus3p, and Ste5-Fus3pp. If the same text-matching wildcard is used multiple times on each side of the equation, then Smoldyn corresponds the first instance in the reactants to the first instance in the products, the second to the second, and so on. For example, if Ste5 can also be phosphorylated, then Ste5* + Fus3* → Ste5*-Fus3* specifies that the binding reaction occurs for all phosphorylation states of both proteins, and that they maintain their phosphorylation states in the reaction. The correspondence can also be given explicitly using the ‘$n’ wildcard on the product side of a reaction, using any value of *n* from 1 to 9, where it represents the *n*’th item of matching text. For example, the previous reaction could also be written as Ste5* + Fus3* → Ste5$1-Fus3$2. Text-matching wildcards in the reactants do not have to appear in the products; for example, Fus3* → X shows that all three Fus3 species decay to the same product. On the other hand, text-matching wildcards in the products must appear in the reactants, meaning that Smoldyn would not accept the reaction X → Fus3*.

Much like the case for species patterns, Smoldyn expands reaction patterns that include structural wildcards to lists of elementary reaction patterns and then performs matching and substitution on these elementary patterns. In the reaction pattern A&* → X*, for example, Smoldyn would first expand it to A* → X* and *A → X*, and would then perform matching and substitution on these two elementary reaction patterns. There are a few possible types of expansions. (*i*) If the reactant and product sides expand to the same number of elementary patterns, then Smoldyn assumes that they correspond to each other sequentially. For example, Smoldyn expands the reaction pattern A|B → C|D to the two reactions A → C and B → D. (*ii*) Smoldyn also accepts patterns that expand to only one elementary pattern on either the reactant or product side, in each case creating a list of reactions that have either the same reactant or product. For example, A|B → X expands to A → X and B → X, and X → A|B expands to X → A and X → B. However, (*iii*) Smoldyn does not accept patterns that expand to different numbers of elementary patterns on the reactant and product sides. For example, Smoldyn rejects the reaction pattern A|B|C → D|E.

In addition to the chemical reaction equation, Smoldyn allows modelers to specify several other reaction parameters. These include the reaction rate constant, how any dissociation products should be arranged, whether molecule serial numbers should be retained, and others. These parameters are entered in the same way for single reactions, reactions defined using wildcards, and reactions defined as rules, described next.

### 3.4 Reaction network expansion

In most cases, Smoldyn acts on input file statements as it encounters them. For example, if Smoldyn encounters a difc statement in an input file, it immediately sets the diffusion coefficient for all species that match to the given species pattern to the given value. Likewise, if Smoldyn encounters a reaction statement, it immediately creates reactions for all currently defined species that match to the given reaction pattern. In this case, Smoldyn issues either a warning or an error if any product names arise that are not currently defined species. Smoldyn does not revisit these statements later on during the simulation.

On the other hand, if the statement is suffixed with “`_rule`”, such as `difc_rule` or `reaction_rule`, then Smoldyn does not act on the statement immediately but instead stores it for future use (after a little preliminary parsing). Smoldyn acts on these statements later on during *rule expansion.* Smoldyn supports two approaches for rule expansion. First, if it encounters an `expand_rules` statement in the input file (followed by “all” or a number), it expands the rules at that point. In this so-called *generate-first* approach (***28***), Smoldyn reads through the rules sequentially and acts on them using the currently defined species. In doing so, if it finds that a reaction specifies a product species that has not been defined, then Smoldyn creates the species. Smoldyn repeats this process for a user-specified number of iterations or until it has fully expanded the reaction network. This generate-first approach is often convenient for small reaction networks because Smoldyn displays all species and reactions before the simulation begins, making it easy to confirm that the network agrees with expectations. Second, the rules can be expanded using the *on-the-fly* approach (***28***), in which Smoldyn acts on the rules at every time step during the simulation, but only as required. In particular, Smoldyn only generates the reactions for a species once the first molecule of the species has actually arisen in the simulation. This prevents the generation of unused species and reactions, which can be a large fraction of the possible ones (***29***). This improves simulation efficiency for large reaction networks and can often enable simulations with infinite reaction networks (see Notes 4-6).

### 3.5 Properties of new species

As mentioned above, each Smoldyn species has several properties, including its diffusion coefficient, graphical display parameters, and set of surface interaction behaviors. Typically, one assigns these properties to all species that are defined in the input file using the `difc, color, display_size, action`, and rate statements, where the last two statements define molecule-surface interaction behaviors. However, if Smoldyn acts on these statements before it performs reaction network expansion (which is always the case for on-the-fly expansion), then they do not apply to any newly generated species. The rule statements described above, such as `difc_rule`, are one way to address this problem. An alternate and often better approach is that Smoldyn can assign species properties automatically by computing reaction product properties from the reactant properties.

It does so using the following assumptions: *(i)* reactants diffuse as though they are roughly spherical, *(ii)* reactant volumes add upon binding, and *(iii)* molecule diffusion coefficients scale as the inverse of the molecule’s radius (***22***). This last assumption follows from the Stokes-Einstein equation, which appears to be reasonably accurate even within cells (***26, 30***). These assumptions lead to the following equations for the product of the generic reaction A + B → AB:

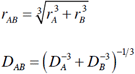

where *r_A_* and *r_B_* are the reactant radii and *D_A_* and *D_B_* are the reactant diffusion coefficients. Smoldyn assigns the product diffusion coefficient as *D_AB_* and computes the product’s graphical display radius from the *r_AB_* equation. Next, Smoldyn computes the product’s display color using a radius-weighted average of the reactant colors. For each of the red, green, and blue colors, it computes the product brightness value using

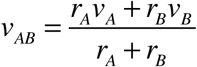

where *v_A_* and *v_B_* are the reactant brightness values. Finally, Smoldyn determines surface interactions for products using the method that the new species behaves like the reactant that has the “greater action,” where the possible actions are ordered with increasing value as: transmission, reflection, absorption, and porting (which is for hybrid simulations). For example, if a surface reflects reactant A and transmits reactant B, then reflection is the greater action, so the surface reflects product AB.

### 3.6 Symmetric species

Reaction networks that include structurally symmetric species often include multiple reactions that form the same products, which increases the effective reaction rate. Consider the A-B-B-A complex for example (see Note 3). It can lose an A monomer from either the left or right sides, whereas the A-B-B complex can only lose an A monomer from the left side, so the former reaction should proceed twice as fast (assuming that all of these A-B bonds are chemically identical). Smoldyn accounts for this by watching for repeated reactions as it expands reaction patterns, and incrementing the associated *reaction multiplicity* when they arise. Smoldyn multiplies the reaction multiplicity by the requested reaction rate constant to compute the total reaction rate constant.

An exception arises to this multiplicity computation if the reaction rule for a bimolecular reaction can match to both possible orderings of a single pair of reactants. For example, the rule * + * → ** can match to the two reactants A and AA as either A + AA or AA + A (see Note 5). Because these two possible reaction orderings typically reflect two different chemical bonds being formed, Smoldyn only considers one of the two orderings (the one in which the reactant’s internal indices are in increasing order).

## 4 Notes

The following notes illustrate the use of wildcards for rule-based modeling using several example problems. These models, and additional files that I used for their analysis, are available in the Smoldyn download package in the subdirectory examples/S94_archive/Andrews_2017. Further information about the models is also available in this paper’s online supplementary information.

### 1 Simple reaction networks with low symmetry

Reaction networks that are conceptually simple and have low symmetry are typically easy to define using wildcards. This is illustrated with an example of second messenger signaling, where extracellular “first messengers” bind to cell receptors, which then release intracellular “second messengers” (***31, 32***). Figure 1A shows a simple model in which a transmembrane receptor (R) can bind an extracellular ligand (L) and/or an intracellular messenger protein (M); a messenger that is bound to a ligand-bound receptor gets phosphorylated (Mp), and phosphorylated messengers lose their phosphates spontaneously (such as from unmodeled phosphatases). The network, which comprises 9 species and 10 reactions (Figure 1B), can be expressed with the following 4 rules using wildcards:

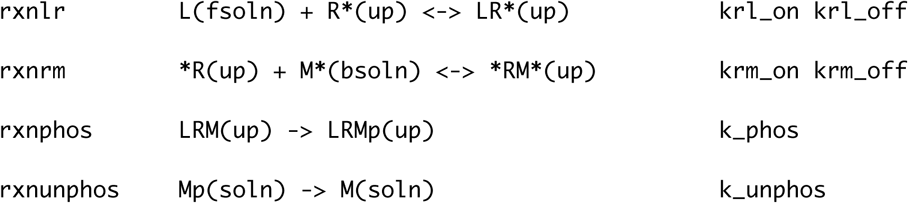

Each line shows the rule name, the reaction rule, and the reaction rate constants. Note that the use of wildcards, which in this case is just the ‘*’ character, enabled each rule to represent a separate process in a clear manner. Also note that the reactant and product states (the spatial localizations given within parentheses) are straightforward to define and reasonably intuitive. Smoldyn uses them to correctly place all receptor complexes at the membrane, ligands in the extracellular space, and messengers in the cytosol (Figure 1C).

**Figure 1.**
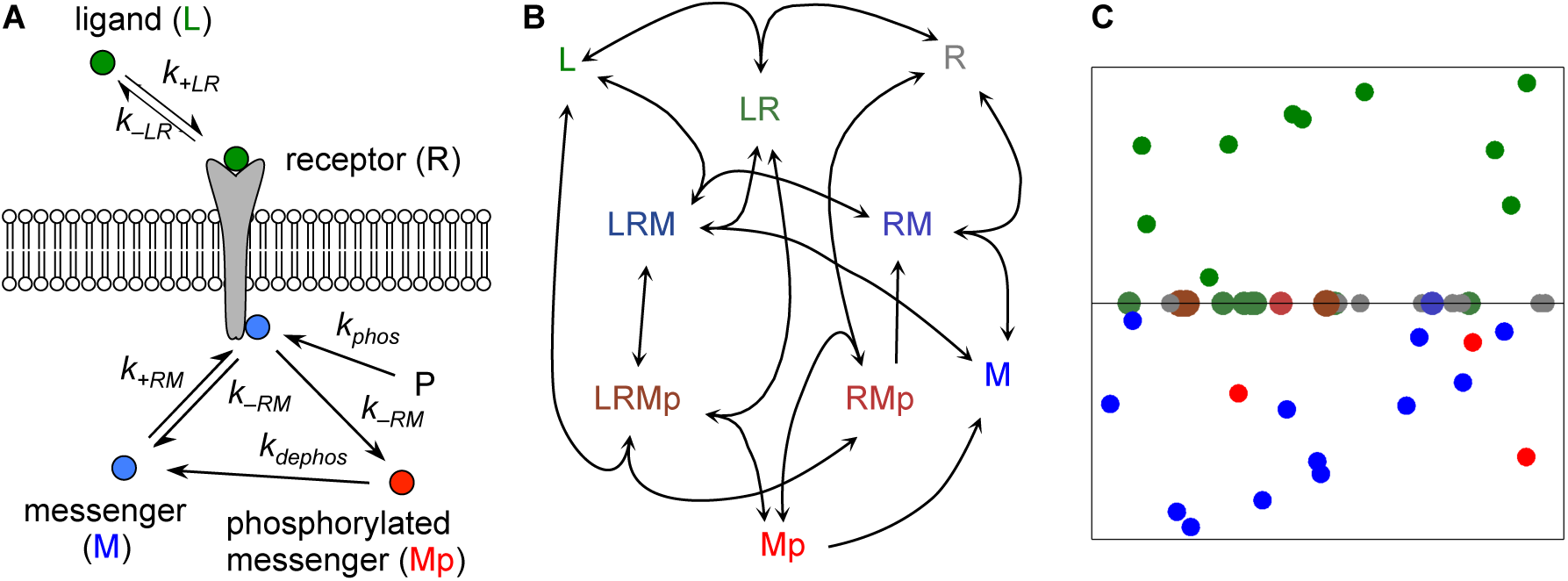
Model of second messenger signaling. (A) Cartoon of the model, showing the components and their interactions. (B) The complete reaction network, where species are shown with the same colors as those generated by Smoldyn. (C) Snapshot of this model simulated in Smoldyn, again using the same color scheme. The line across the middle represents the membrane, the region above the line is the extracellular region, and the region below the line is the cytoplasm.

### 2 More complicated networks with low symmetry

Figure 2A shows a slightly more complicated example, but one that still includes asymmetric complexes. It shows a model of transposon excision that was developed to answer the question of how DNA transposons regulate their copy numbers so that they do not overproduce themselves and then kill their hosts ***(33)*** (transposons are mobile sections of DNA that can be amplified as they move from one location in the genome to another). In the model, the A-B species is a transposon with ends ‘A’ and ‘B’, and T_2_ is a transposase dimer, an enzyme that binds to and cuts transposon ends. The transposase can be non-specifically bound to DNA (T_2,nsb_) or freely diffusing in the nucleus (T_2_). At low transposase concentrations: a T_2_ binds to a transposon end to form a singly-bound transposon (T_2_A-B or A-BT_2_), this DNA forms a loop, the same T_2_ binds to the other transposon end (AT_2_B), and the transposase cuts out the transposon (the reaction with rate *k_3_*). In the model, the transposition products conserve the reactant amounts and create an X molecule as a transposition counter although, in actuality, transpositions can produce additional transposons, amplifying the transposon in the genome. At high T_2_ concentrations: transposases bind to singly-bound transposons to create doubly-bound transposons (T_2_A– BT_2_), which cannot undergo transposition, thereby regulating the process. This model can be expressed using wildcards as

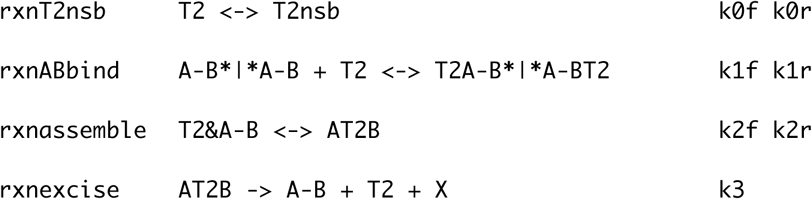

The OR operators in rxnABbind indicate that T_2_ can bind to either the left of A-B* (A-B or A-BT_2_) or the right of *A-B (A-B or T_2_A-B). The permutation operator in rxnassemble indicates that both T_2_A-B and A-BT_2_ react to form AT_2_B.

**Figure 2.**
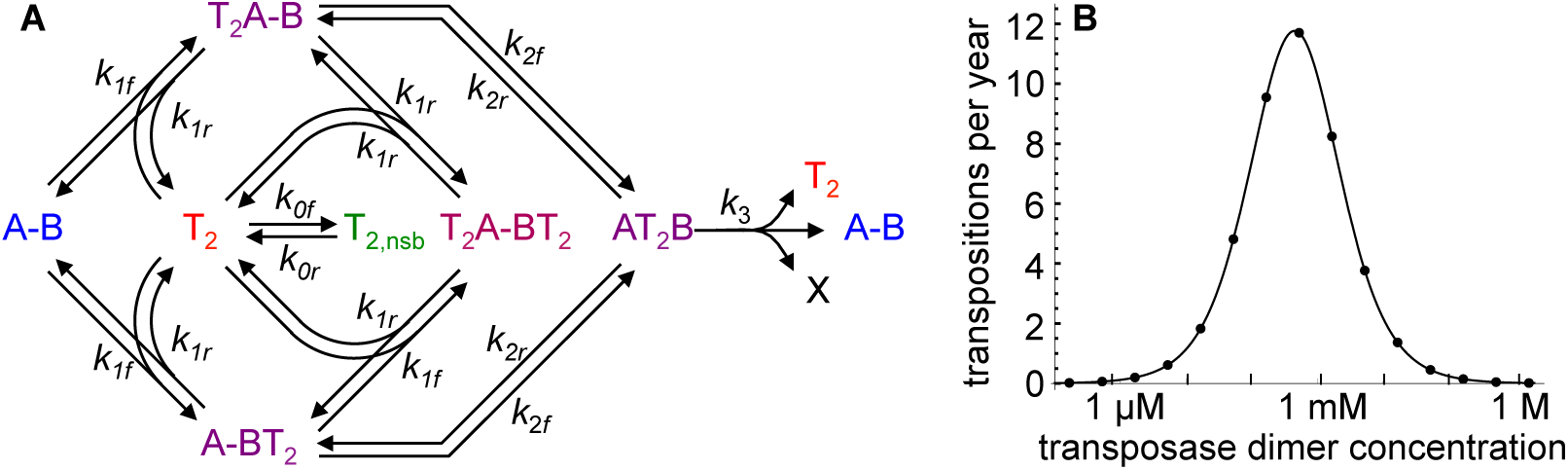
Model of transposase dynamics modified from ref. ***(33)***. (A) Reaction network, where A-B is a transposon and T_2_ is a transposase dimer. Colors are those generated by Smoldyn. (B) Transposition rate for a single transposon as a function of the total transposase dimer concentration within the nucleus. Points represent simulation data generated with ordinary differential equations and simulated in Mathematica and the line represents an analytical theory for the transposition rate (supplementary information).

Expanding these reaction rules with Smoldyn produced the reaction network shown in Figure 2A, as anticipated. The physiological rate constants (***33***) vary extremely widely (e.g. *k_0f_*= 10^5^ s^-1^ and *k_2f_* = 4.3×10^-4^ s^-1^), meaning that Smoldyn would have to use short time steps to resolve the fast reactions but also run for a very long time to observe the slow reactions, so I simulated these reactions deterministically instead using Mathematica. Figure 2B compares the simulated transposition rates with values derived from analytical theory (supplementary information), showing excellent agreement.

### 3 Symmetric complexes, modeled with asymmetric notation

Reaction networks that include structurally symmetric protein complexes, such as dimers and higher oligomers (***34***), generally require a little more care. In particular, it is often the case that a single complex can be represented correctly in multiple ways, leading to the question of whether the model notation should just include one of the ways, or all of them. Which approach is simplest depends on the specific problem; this note shows an example of the former approach, in which each complex is represented in just one way.

Figure 3A shows a simple model of reversible dimer assembly for a symmetric complex that has the form A-B-B-A, a form that is loosely based upon receptor tyrosine kinases such as the epidermal growth factor and insulin receptors (***35***). The model includes the monomers A and B, dimers AB and BB, trimer ABB, and the tetramer ABBA. The notation is asymmetric in that it includes the species AB but not the species BA, which would be chemically identical. Similarly, it includes ABB but not BBA. It can be expressed with the reaction rules:

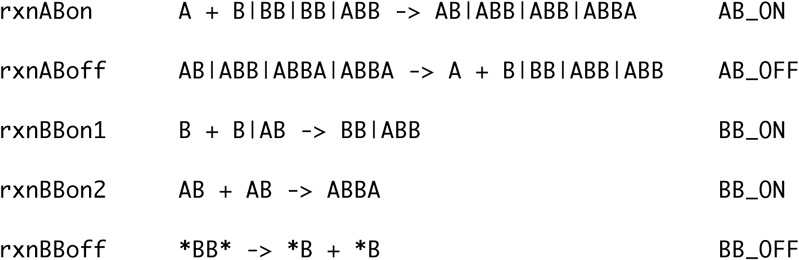

These rules make heavy use of the OR operator. For example, the first reaction rule shows that A can bind to any of B, BB, BB, or ABB, and the products are, respectively, AB, ABB, ABB, and ABBA. The repeated BB reactants in this rule reflect the fact that A can bind to either the left or right side of BB so the rate constant for this reaction should be twice the listed value (`AB_ON`). Similarly, in the second reaction rule, ABBA dissociates twice to A + ABB to reflect the two A-B bonds in ABBA. These rules are somewhat inelegant in that they do not reflect the symmetry of the system, include strings of OR operators, and only include irreversible reactions despite the fact that the model reactions are reversible. This inelegance arises from the decision to use asymmetric notation and from limitations in the wildcard approach. Nevertheless, these rules are substantially simpler than the full list of 12 reactions.

**Figure 3.**
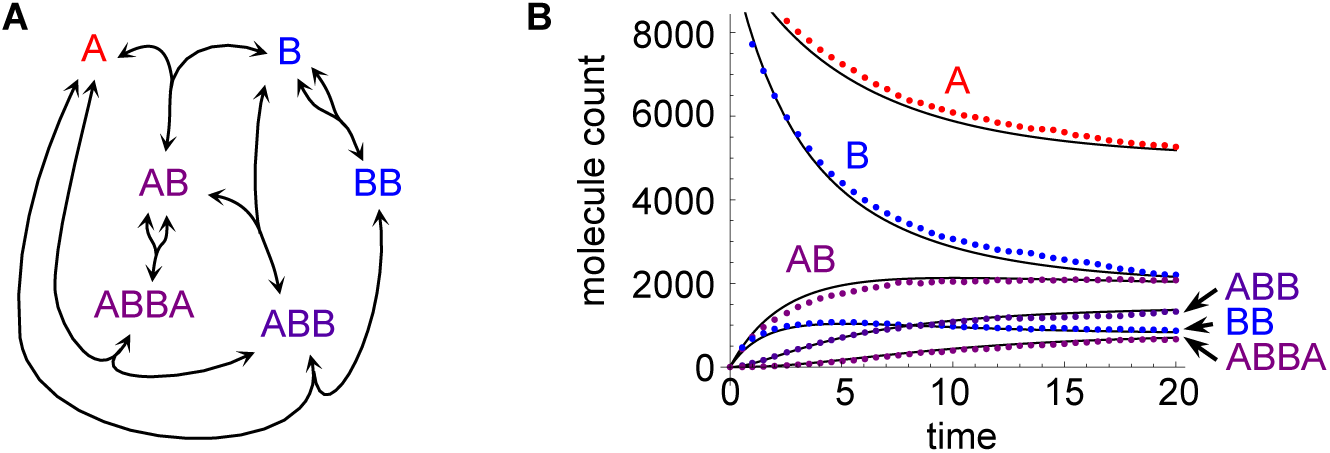
Model of symmetric complexation using asymmetric notation. (A) Reaction network for binding between A and B components that can assemble into the A-B-B-A complex. (B) Black lines show reaction kinetics computed from manual reaction network expansion and simulated with ordinary differential equations using Mathematica; colored points show reaction kinetics from Smoldyn’s expansion of wildcard rules and then simulation. Colors in both panels are those generated by Smoldyn. Simulation parameters: AB_ON = 10, AB_OFF = 0.05, BB_ON = 8, BB_OFF = 0.03, 10,000 initial A molecules, 10,000 initial B molecules, volume of 100^3^, time from 0 to 20 with steps, in Smoldyn simulation, of 0.05.

The reaction network that Smoldyn computed from these rules was identical to ones that arose from BioNetGen and manual expansion (***22***), validating the rule approach. Figure 3B shows that a Smoldyn simulation that was defined with these rules agreed well with a deterministic simulation of the same network, computed using ordinary differential equations.

### 4 Symmetric complexes, modeled with symmetric notation

This note continues on the topic of symmetric complexes, but now using symmetric notation. Here, if a complex can be represented correctly in multiple ways, the approach is to not just pick one of them but to use all of them. This increases network complexity due to the greater number of species and reactions, but can simplify the reaction rules through maintenance of the network symmetry.

*E. coli* bacteria have several mechanisms for locating their cell division plane at the cell center, one of which is to prevent division elsewhere with the Min system (***36,37***). Here, the combined actions of the MinD and MinE proteins create a spatiotemporal oscillation between the cell poles that keeps the co-localized MinC away from the cell center; MinC inhibits division apparatus formation, thus inhibiting cell division away from the cell center. This system has been modeled extensively (***38,39***) but few models explicitly represent MinD or MinE dimerization (***40***), despite the fact that both have dissociation constants that are comparable to their intracellular concentrations (***41,42***). Interestingly, MinD only dimerizes when bound to ATP (***43***) and MinD only hydrolyzes ATP when it is dimeric (***44***).

Figure 4A shows a model of MinD nucleotide binding and dimerization. All species are MinD proteins, but bound to different co-factors: ‘T’ represents MinD bound to ATP, ‘D’ represents MinD bound to ADP, and ‘A’ represents MinD bound to neither (‘A’ stands for apo). Pairs of these symbols, such as ‘TT’, represent dimers. Three of the dimers are heterodimers that the model represents using both possible orderings, such as DT and TD. The model can be described with the following rules, of which the first three represent nucleotide substitution, and subsequent ones represent dimerization, dimer dissociation, and ATP hydrolysis:

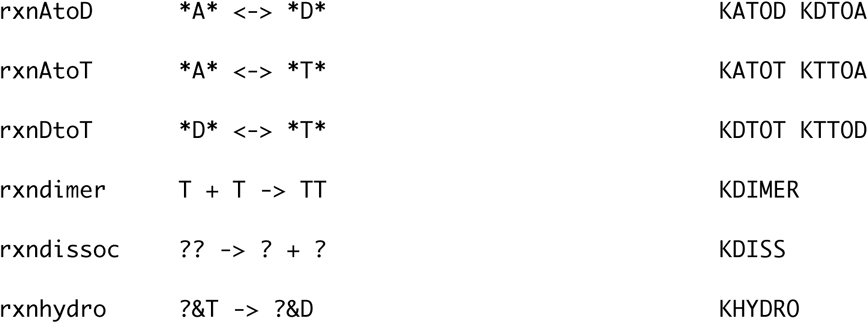

Maintaining the reaction network symmetry in the model notation enabled simple and elegant reaction rules in this case. Note the use of the ‘?’ wildcard; for example, rxndissoc uses it to indicate that all dimers dissociate with the same rate constant. Also, use of the permutation operator in the last rule shows that any dimer with a ‘T’ in it, regardless of whether the ‘T’ is the first or second symbol, is able to perform hydrolysis:

**Figure 4.**
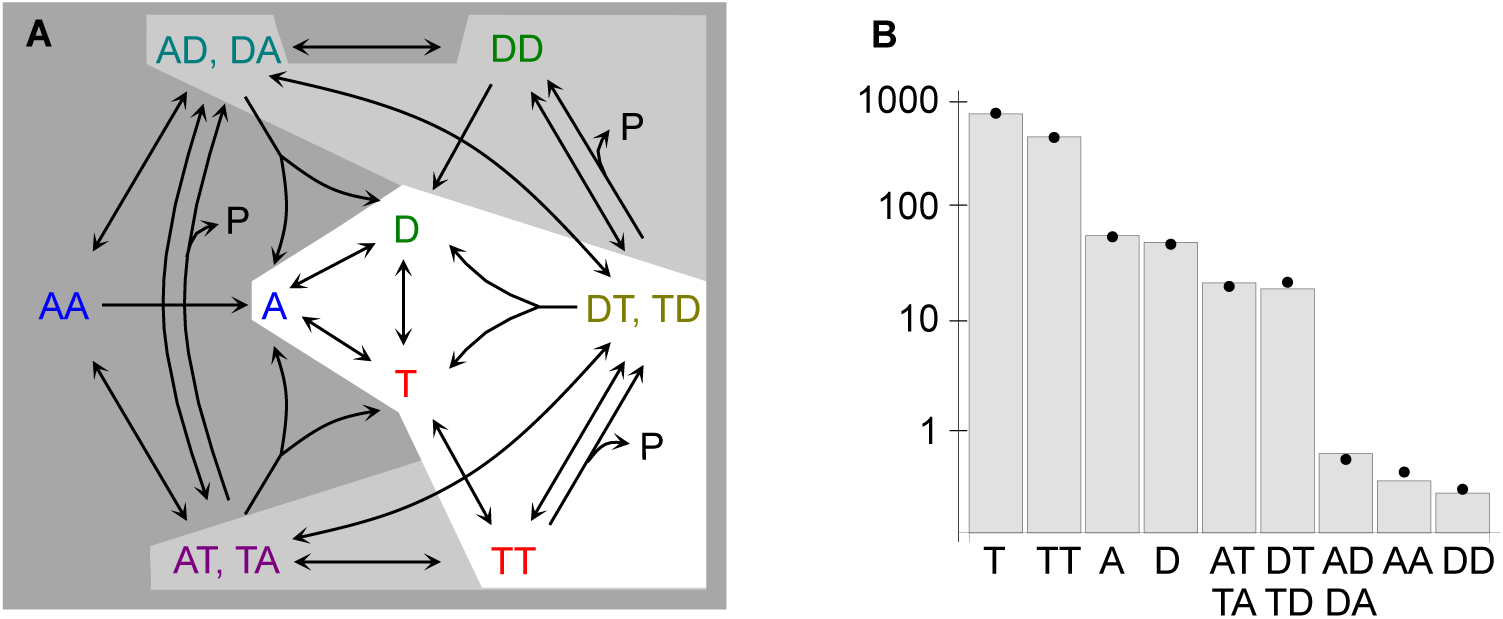
Model of *E. coli* MinD dimerization and nucleotide binding. (A) Reaction network. Background shading illustrates on-the-fly simulation for a simulation in which A, D, T, TT, and either DT or TD have arisen. White regions are explored, light grey are generated by not explored, and dark grey are not generated; see the main text. (B) Species abundance in a single cell at steady-state using physiologically reasonable parameter estimates. Bars are deterministic values computed by simulating the network in Mathematica using ordinary differential equations and points are averages of Smoldyn simulation values (*n* = 20). Parameters are listed in the supplementary information.

Figure 4A also illustrates on-the-fly network generation for this model using background shading. It depicts the situation in which the only species that have arisen in the simulation so far are A, D, T, TT, and DT and/or TD. They are over a white background to show that this region of the network has been explored. Species and reactions in the adjacent light grey regions have been generated by Smoldyn so that they could be used, but they have not actually been used in the simulation so far. Species and reactions in the dark grey regions have not been generated yet (and may not require generation), which saves computation and computer memory.

Figure 4B shows the number of molecules of each species at steady-state, where the bars are from a deterministic simulation in Mathematica and the points are average values from a Smoldyn simulation. It shows that most of the MinD is bound to ATP and is either monomeric or dimeric. These results were computed from physiologically reasonable parameters for a single cell (supplementary information), but do not account for membrane or MinE interactions.

### 5 Polymerization with identical monomers

Cellular polymers include (*i*) microtubules and actin, which are important for cell structure and intracellular transport, (*ii*) intermediate filaments, which provide mechanical strength, (*iii*) DNA and RNA, which encode genetic information, (*iv*) polysaccharides, which provide structure and store energy, and sometimes (*v*) amyloid fibrils, which can cause neurodegenerative diseases (***45-47***). Most of these polymers assemble at one or both ends, although some can also anneal, meaning that two polymers join end-to-end.

Figure 5A shows a polymer model that assembles and disassembles at one end (called “polymer_end1”). It can be expressed with the reaction rule

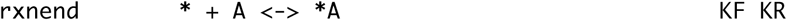

where ‘A’ is a single polymer unit and KF and KR are the forward and reverse reaction rate constants. The isolated asterisk was adequate in this rule because this model did not include other species, but would have created unintended reactions otherwise. A simulation that started with 20,000 monomers and used on-the-fly expansion showed that the polymers exhibited an exponential length distribution at equilibrium, in agreement with theory (***48***) (Figure 5C, supplementary information). On completion, this model had 40 species and 77 reactions.

**Figure 5.**
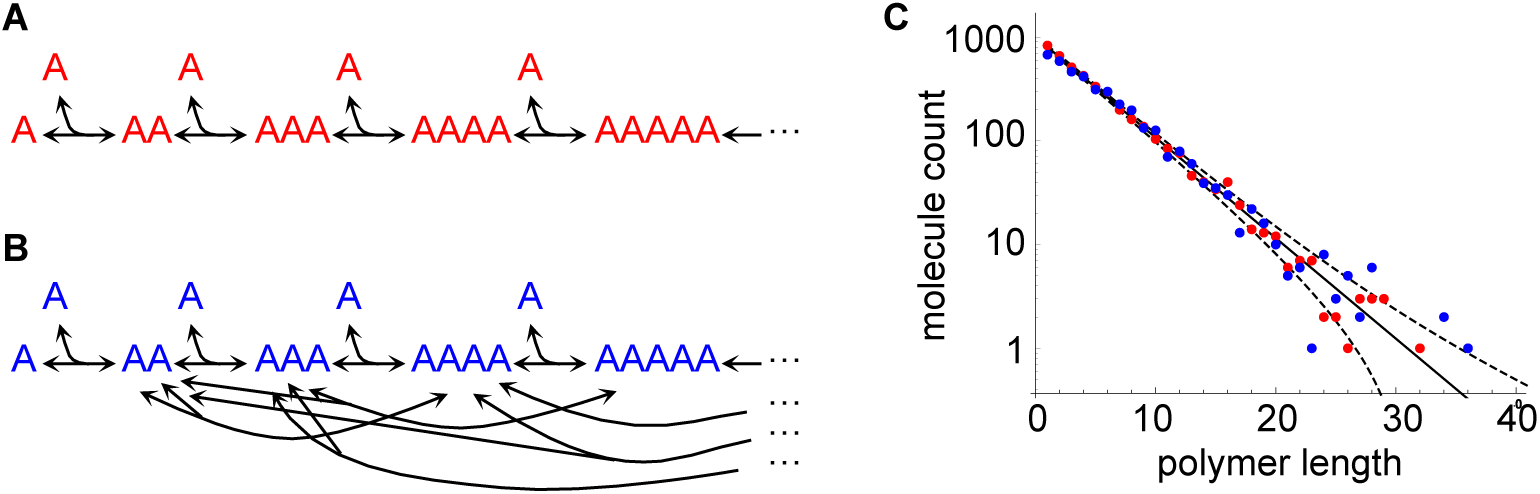
Models of polymerization. (A) Reaction network for polymers that can add or lose units from a single end. (B) Part of a reaction network for polymers that can add or lose units from their ends, and can also break and anneal. (C) Equilibrium length distributions of polymers from a simulation of the end-polymerization model (“polymer_end1” model, red points), a simulation of the breaking and annealing model (“polymer_mid” model, blue points), and analytical theory (solid black line). Dashed lines show the theoretical standard deviations.

Some limitations of the method were interesting. (*i*) This simulation represented polymer lengths by listing their units rather than with numbers (e.g “AAA” rather than “A3”), so polymers were limited to 256 units because that is the longest species name that Smoldyn allows. (*ii*) Smoldyn represents these polymers as spheres rather than as extended filaments; this is clearly inaccurate for stiff polymers, although actually reasonably accurate for highly flexible polymers which tend to collapse into loose clusters (***48***). In the latter case, Smoldyn computes polymer radii as increasing as *L*^1/3^, where *L* is the polymer length, whereas the ideal scaling for freely jointed chains is *L*^1/2^ (***47,49***). And (*iii*) Smoldyn computes the polymer diffusion coefficients as decreasing as *L*^-1/3^, as compared to the *L*^-0.6^ scaling that is typically observed experimentally for polymers (***47***).

A more serious flaw with this model is that Smoldyn assigns the same reaction rate to all association reactions. This is the correct behavior for the given reaction rule, but does not account for the fact that the A + A → AA reaction can happen in either of two ways: either of the two reactant monomers can end up at the “left” end of the product. The following reaction rules (model “polymer_end2”) fix this flaw

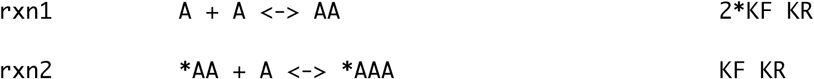

Here, monomer association proceeds twice as fast as association of higher polymers. Results from these latter rules agreed with a comparable model written in BNGL (***22***) and, at equilibrium, they also showed an exponential length distribution for all polymers with more than one monomer (see supplementary information).

Figure 5B shows a similar model, but one which represents polymers that can anneal and break (model “polymer_mid”). It can be expressed with the reaction rule

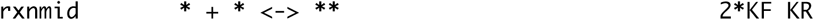

As above, the association reaction rate was doubled to account for the fact that either of the two reactants can end up on the “left” side of the product. This follows from the fact that Smoldyn only considers a single ordering for any particular pair of reactants; for example, it generates the reaction A + AA → AAA but not also AA + A → AAA. This model reached equilibrium much faster than the former ones but produced essentially the same exponential length distribution as the “polymer_end1” model (Figure 5C). This model led to a much larger reaction network, with 151 species and 4037 reactions, because each species can participate in many more reactions.

### 6 Polymer sequences and chemical structures

The pattern matching aspects of the wildcard method enable it to be used to define reactions that are specific to individual polymer sequences and chemical structures. These applications would be possible using BioNetGen or other rule-based modeling approaches, but would generally not be as convenient.

The central dogma of molecular biology is that cells transcribe DNA to mRNA and then translate mRNA to protein (***45***). Figure 6A shows that this process can be modeled using wildcards if sequences are reasonably short. The reaction rule

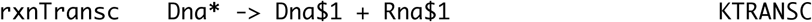

performs transcription, where “Dna” and “Rna” are prefixes that indicate the sequence type and the “$1” portions of the products show that the same text gets substituted into each one. Ideally, this rule would not only preserve the sequence, which it does, but also replace all T symbols, for DNA thymine bases, with U symbols, for RNA uracil bases. However, there is no easy way to do so with the wildcard method as it is currently designed. A wildcard approach that used regular expressions, which are more sophisticated pattern matching approaches, would solve this problem but would also be more difficult to use. The following reaction rules perform translation by modeling ribosome (“Rib”) binding to the beginning of an mRNA sequence, translation of each codon, and finally dissociation of RNA, ribosome, and protein (“Prot” prefix).

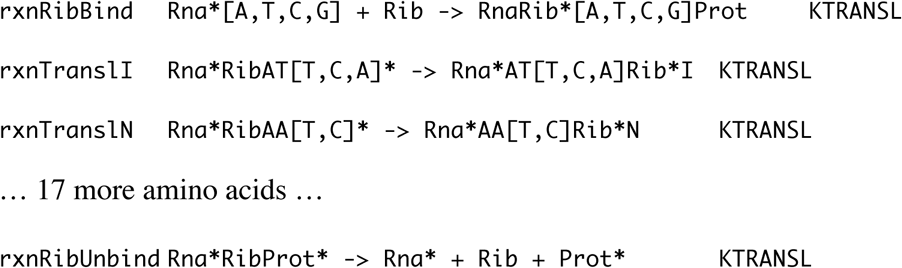

In `rxnTransl`I reaction rule, for example, any of the RNA codons ATT, ATC, and ATA (using T instead of U) code for isoleucine, so the product shows that the ribosome moves forward by three basepairs and an ‘I’, for isoleucine, is appended to the growing protein. Three final reaction rules encode for DNA mutations and RNA and protein degradation, respectively,

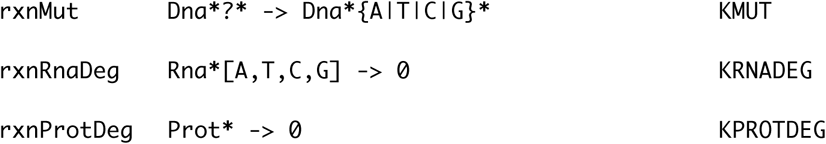

Figure 6B shows results from a simulation of this model that started with one DNA molecule, DnaATCAATATT. Initially, it was transcribed to RnaATCAATATT and then translated over multiple steps to ProtINI (isoleucine-asparagine-isoleucine). At a simulation time of about 1.9 hours, the DNA mutated, leading to a slightly different RNA sequence and production of protein IYI. The protein molecule counts show a large variation because they amplify the RNA counts, which have high variation due to their low copy numbers (***50***)

**Figure 6.**
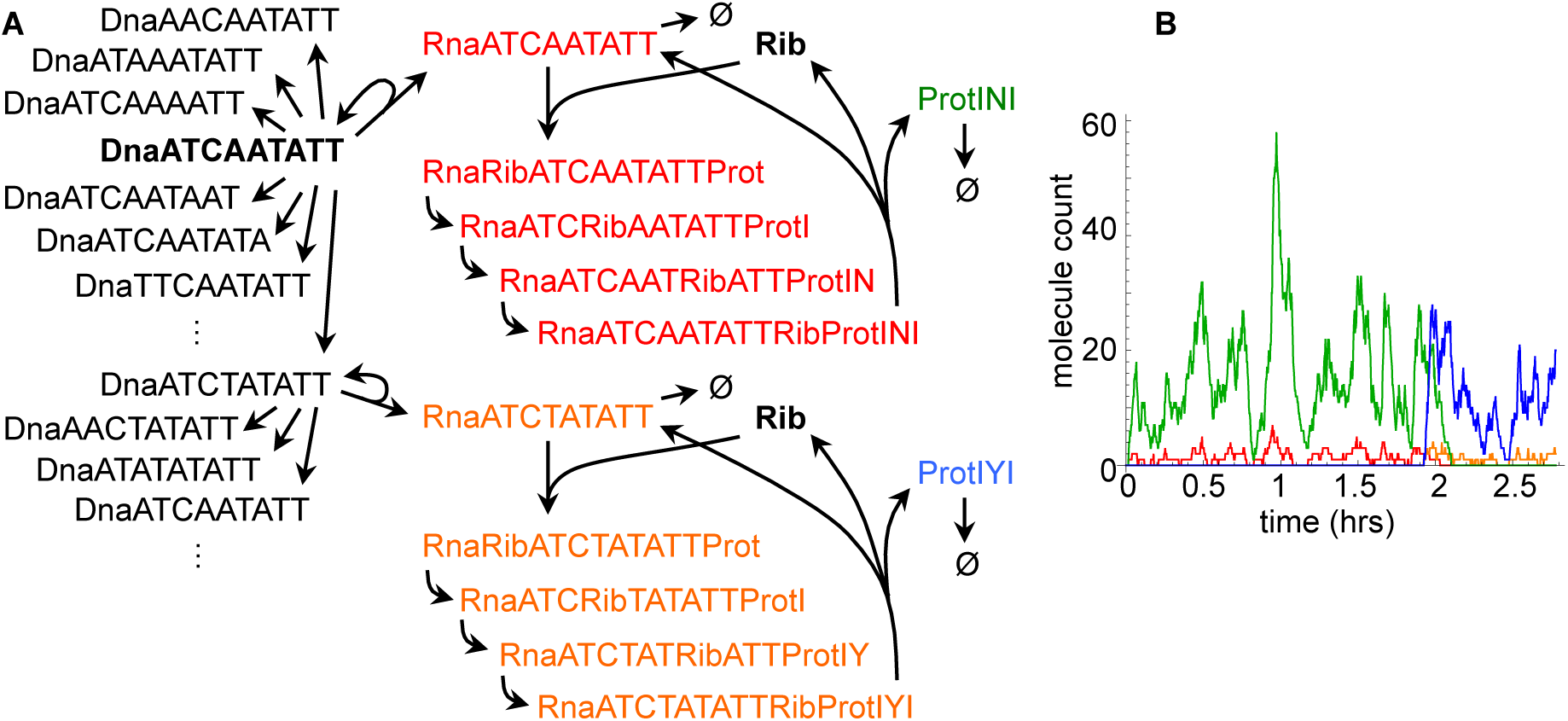
Model of DNA transcription and then RNA translation. (A) Diagram of the model, showing some of the species that arose during a particular simulation run that used on-the-fly network expansion. The starting DNA sequence is shown in black and bold face. It could mutate to other sequences, shown above and below in black. It was also transcribed to RNA, shown in red, and these RNAs were translated one codon at a time to produce the polypeptide ProtINI, shown in green. Mutation actually happened at simulation time of about 1.9 hours, at which point the new DNA was transcribed to the RNA shown in orange and it was translated to the protein shown in blue. (B) Copy numbers of the RNA and protein molecules from the same simulation, using the same colors as panel A.

Wildcards can also be used to define reactions based on chemical structures that are not sequence data. In particular, they are useful in conjunction with the SMILES notation (***51***), a scheme that allows most chemical complexes to be uniquely expressed using a single line of normal text characters (e.g. ethanol, CH_3_CH_2_OH is CCO in SMILES notation). As an example, the *E. coli* lipid synthesis pathway includes several enzymes that act repeatedly on lipids, adding a two carbon groups with each repetition (***52***). Each enzyme is specific to a particular chemical functional group but has low specificity with regard to the lipid chain length. This can be representing using wildcards starting with the 10-carbon lipid cis-3-decenoyl-ACP, written in SMILES notation as (***53***)C(=O)C/C=C\CCCCCC. Here, ACP is an abbreviation for acyl carrier protein, the C(=O) portion represents a carbonyl group, the /C=C\ portion represents a cis-conformation double bond, and the CCCCCC portion represents a saturated hydrocarbon tail. The reaction rules are

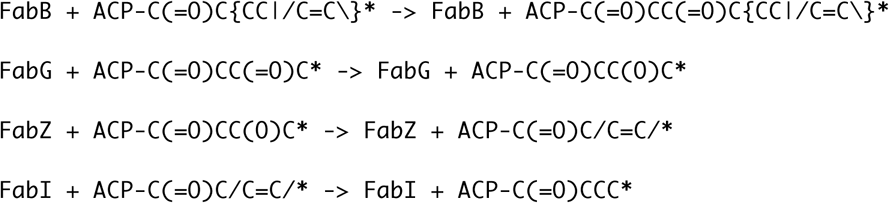

In the first reaction, FabB adds a carbonyl and extra carbon, C(=O)C, to the chain. Next, FabG reduces the newly added carbonyl to a hydroxyl, C(O); FabZ reduces the hydroxyl to a trans-conformation carbon-carbon double bond, /C=C/; and then FabI reduces the double bond to a single bond, CC. The end result is that the cis-3-decenoyl-ACP gets lengthened by two carbons to cis-3-dodecenoyl-ACP. Application of these rules to this longer lipid adds yet more carbons.

Both the nucleic acid sequence model and this lipid synthesis model would undoubtedly be simpler and more generalizable if they were developed using software designed specifically for the tasks. However, the fact that they can be developed using wildcards shows the method’s versatility.

## Acknowledgement

I thank Ronnie Chalmers, Akintunde Emiola, Jim Faeder, and Karen Lipkow for useful discussions. Much of this work was carried out during a visit to the Isaac Newton Institute for Mathematical Sciences, for which I thank Radek Erban, David Holcman, Sam Isaacson, and Konstantinos Zygalakis, who were the program organizers, and the Institute Staff. I also thank Roger Brent, Erick Matsen, and Harlan Robbins for providing space for me at the FHCRC, where the work was completed. This work was supported by a Simons Foundation grant awarded to SSA and by EPSRC grant EP/K032208/1 awarded to the Isaac Newton Institute.

## Supplementary Information

### 1 Second messenger signaling

#### Smoldyn input file

~~~
# Smoldyn configuration file to test wildcards in reactions
# This file simulates second messenger signaling with ligand (L), receptor (R),
# and messenger (M). R is membrane-bound and can bind L and/or M. If it binds
# both, then M gets phosphorylated to Mp.

define krl_on  20
define krl_off 0.005
define krm_on  10
define krm_off 0.1
define k_phos  1
define k_unphos        0.01

# Graphical output
graphics opengl_good

# System space and time definitions
dim 2
boundaries x 0 100
boundaries y 0 100
time_start 0
time_stop 1000
time_step 0.05

# Molecular species and their properties
species L R M Mp
difc L(all) 3
difc R(up) 0.2
difc M(all) 2
difc Mp(all) 1.5
color L(all) green
color R(all) grey
color M(all) blue
color Mp(all) red
display_size all(all) 2

# Reactions
reaction_rule rxnlr L(fsoln) + R*(up) <-> LR*(up) krl_on krl_off
reaction_rule rxnrm *R(up) + M*(bsoln) <-> *RM*(up) krm_on krm_off
reaction_rule rxnphos LRM(up) -> LRMp(up) k_phos
reaction_rule rxnunphos *Mp(soln) -> *M(soln) k_unphos

expand_rules all

# Surface parameters
start_surface membrane
action all(all) both reflect
panel rect +1 0 50 100
end_surface

start_surface outsides
action all(all) both reflect
panel rect +x 0 0 100
panel rect -x 100 0 100
panel rect +y 0 0 100
panel rect -y 0 100 100
end_surface

# initial molecules
surface_mol 20 R(up) membrane all all
mol 20 L 50 80
mol 20 M 50 20

#text_display time Mp M* LR*(all)

end_file
~~~

### 2 Transposon dynamics

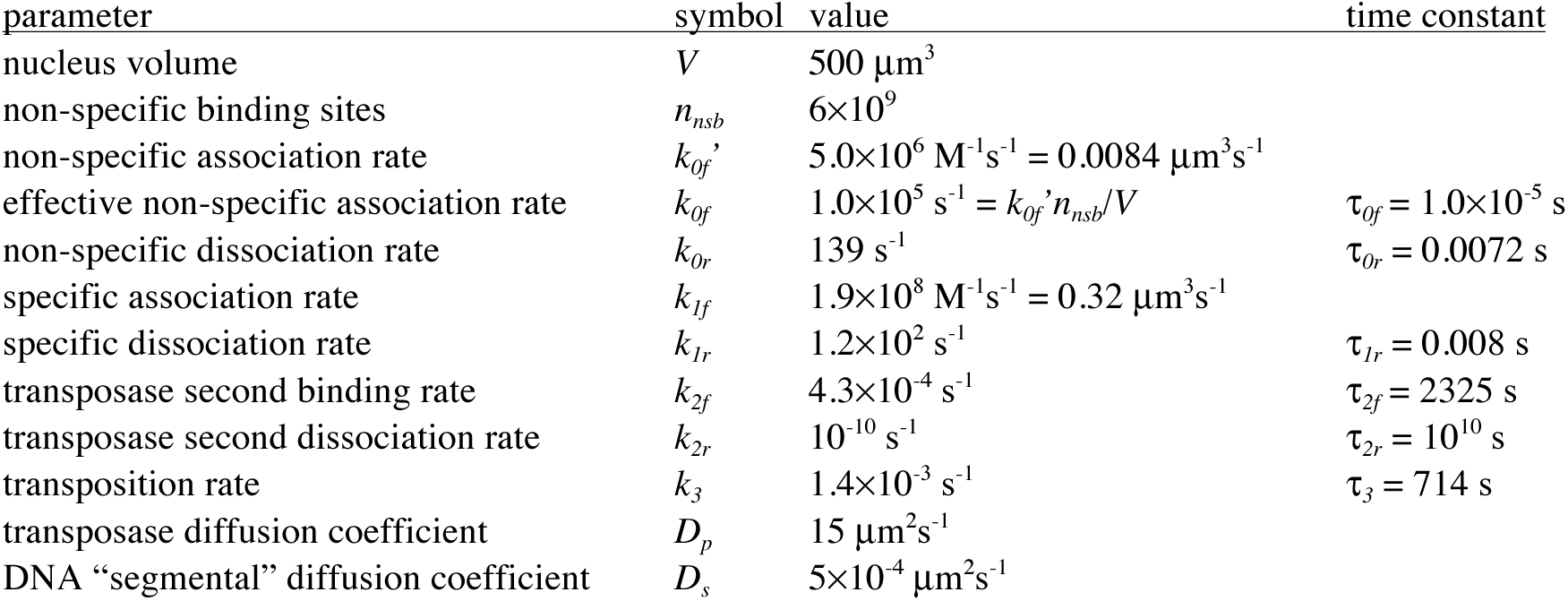
Model parameters. Model parameters were the values determined for the *in vivo* situation from ref. ***(1).*** They are:

#### Analytical model

The above table shows that the reaction time constants, which are the inverse of the reaction rate constants, vary over 15 orders of magnitude. This suggests the possibility of simplifying the model by separating the fast processes from the slow ones, and treating the fast ones as though they are at equilibrium. Indeed, this approach works well here because reactions 0 and 1, which interconnect the group of species T_2,nsb_, T_2_, A-B, T_2_A-B, A-BT_2_, and T_2_A-BT_2_, are much faster than reaction 2, which leads out of this group, to AT_2_B. Thus, assume that this group of species is at equilibrium.

Focus first on reaction 0. At equilibrium, the concentration ratio between T_2_ and T_2,nsb_ is

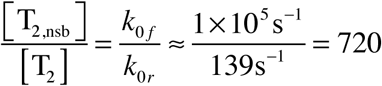

This shows that 720 times as much transposase is non-specifically bound than is freely diffusing. With the assumption that there are many fewer transposons than total transposase dimers, then an insignificant number of transposases will be bound to transposons. Define [T_2,total_] as the sum of the non-specifically bound and freely diffusing transposase dimer concentrations, [T_2,nsb_] + [T_2_], which is a close approximation to the actual total transposase dimer concentration. Using this, the fraction of total transposase that is freely diffusing can be approximated as

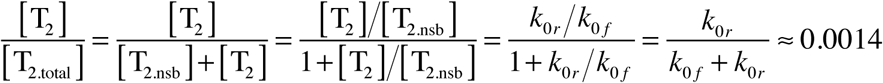

Focus next on reaction 1. At equilibrium, the following concentration ratios are equal to each other and can be given in terms of *k_1f_* and *k_1r_,*

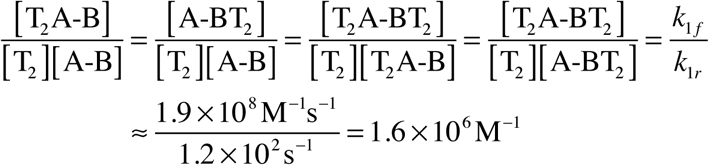

Define the association constant *K* as

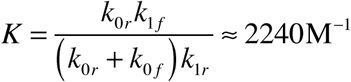

The inverse of *K*, which is about 500 μΜ, is a dissociation constant equal to [A-B][T_2_,_total_]/[T_2_A-b]; it is the total transposase dimer concentration at which there are equal concentrations of transposon in the unbound state A-B as in the singly bound state T_2_A-B. Also define the transposon concentration sum as

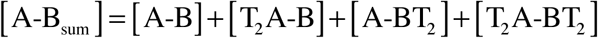

This is close to the total transposon concentration, except that it ignores the concentration of transposons in the AT_2_B state. These are not necessarily negligible because reaction 3 is slow. The fraction of transposons in each binding state can be computed by combining the above equations, giving

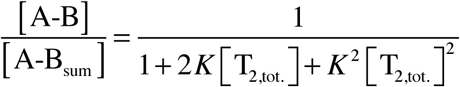

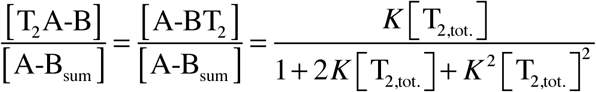

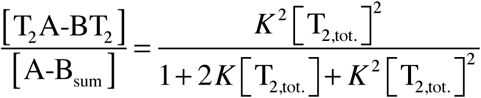

Finally focus on reactions 2 and 3. The concentration of AT_2_B can be computed by setting its rate equation to 0,

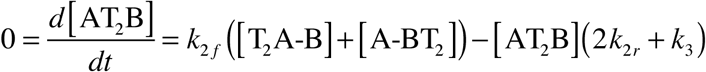

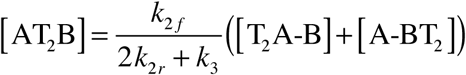

If *k*_3_ >> *k_2f_,* then this concentration is negligible when compared to the total transposon concentration. However, this condition does not hold well for the parameters given above, where *k_3_* is only about 3 times larger than *k_2f_.* Thus, the total transposon concentration needs to be computed. Defining it as [A-B_tot_], it is [A-B_sum_] + [AT_2_B]. So, add [A-B_sum_] to each side of the above equation, substitute in for the singly-bound transposon concentrations, and simplify, to get

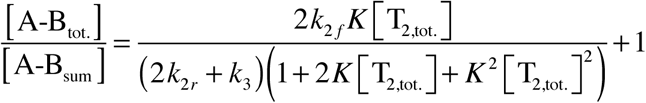

Next, use the fact that reaction 2 is upstream of reaction 3 and has a negligible reverse reaction rate, so its rate is the transposition rate. In other words, any transposon that undergoes reaction 2 will almost certainly go on to transposition, so the rate of reaction 2 is the rate of transposition. Define the rate of reaction 2 as *ϕ*, for the transposition flux, which is

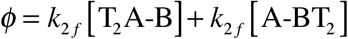

Substituting in the above values gives

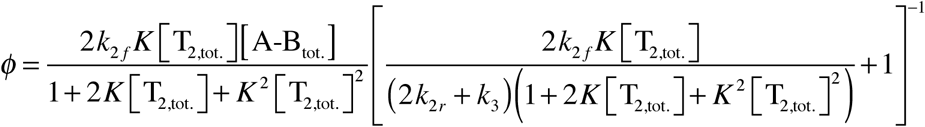

The final term can be neglected if *k*_3_ >> *k_2f_.* This final result is the transposition rate as a function of the transposase dimer concentration. For the most part, it depends only on the rate of DNA ring closing, which is *k*_*2f*_, and on the association constant for transposase onto transposons, which is *K.*

#### Smoldyn input file

This file was only used to verify that the reaction network that was defined using wildcards exactly matched the one that was created manually. The Smoldyn simulation proceeded too slowly for it to be practical.

~~~
# Smoldyn configuration file for transposon dynamics
# Units are microns and seconds

define k0f 1e5
define k0r 139
define k1f 0.32
define k1r 1.2e2
define k2f 4.3e-4
define k2r 1e-10
define k3  1.4e-3
define Dp  15
define Ds  5e-4

define LEN 500^(1/3)

# Graphical output
graphics opengl
graphic_iter 100

# System space and time definitions
dim 3
boundaries x 0 LEN p
boundaries y 0 LEN p
boundaries z 0 LEN p

time_start 0
time_stop 100
time_step 0.0001

# Molecular species and their properties
species A-B T2 T2nsb

difc A-B      Ds
difc T2       Dp
color A-B     blue
color T2      red
color T2nsb   green

display_size all 2

# Reactions
Reaction rxnT2nsb T2 <-> T2nsb k0f k0r
reaction_rule rxnABbind A-B*I*A-B + T2 <-> T2A-B*I*A-BT2 klf klr
reaction_rule rxnassemble T2&A-B <-> AT2B k2f k2r
reaction_rule rxnexcise AT2B -> A-B + T2 + X k3

expand_rules all

# initial molecules
mol 1 A-B LEN/2 LEN/2 LEN/2
mol 10 T2 u u u

#output_files transposaseout.txt
#output_format CSV
#cmd B molcountheader transposaseout.txt
#cmd N 100 molcount transposaseout.txt

end_file
~~~

### 3 Complexation of ABBA structure

#### Smoldyn input file

~~~
# Smoldyn configuration file to test wildcards in reactions
# This file produces a multimeric complex with structure A-B-B-A

define AB_ON   10
define AB_OFF  0.05
define BB_ON   8
define BB_OFF  0.03
define A0      10000
define B0      10000

# Graphical output
graphics opengl_good

# System space and time definitions
dim 3
boundaries x 0 100 p
boundaries y 0 100 p
boundaries z 0 100 p
time_start 0
time_stop 20
time_step 0.05

# Molecular species and their properties
species A B
difc A 2
difc B 1
color A red
color B blue
display_size A 1
display_size B 1

# Reactions
reaction_rule ABon A + B|BB|BB|ABB -> AB|ABB|ABB|ABBA AB_ON
reaction_rule ABoff AB|ABB|ABBA|ABBA -> A + B|BB|ABB|ABB AB_OFF
reaction_rule BBon1 B + B|AB -> BB|ABB BB_ON
reaction_rule BBon2 AB + AB -> ABBA BB_ON
reaction_rule BBoff *BB* -> *B + *B BB_OFF

expand_rules all

# initial molecules
mol A0 A u u u
mol B0 B u u u

output_files abbaout.txt
output_format csv
cmd i 0 20 0.5 molcount abbaout.txt

#text_display time A B AB BB ABB ABBA

end_file
~~~

### 4 MinD dimerization

#### Model parameters

- Cell volume. The cytoplasmic volume of an *E. coli* cell is roughly 0.67 μm^3^ (from BioNumbers ID 100011 (***2***) and (***3***)). However, this model assumes a slightly larger volume of 1 μm^3^, which is 1 fl, in part because the MinD system is of particular interest in cells that are about to divide, which are larger than average cells.
- Number of MinD proteins. Shih et al. reported that there are about 2000 MinD protein copies per cell (***4***).
- ATP concentration and molecules. The ATP concentration varies widely between different individual cells but has an average value of about 1.54 mM (from BioNumbers ID 111006 (***2***) and (***5***)). In a volume of 1 fl, this is 930,000 molecules.
- ADP concentration and molecules. The ratio between ATP and ADP in exponentially growing cells ranges from 5.6 to 10.3, depending on growth conditions (BioNumbers ID 103384 (***2***) and (***6***)). This model assumes a ratio of 8, using the middle of this range. With this ratio and the assumption of 930,000 ATP molecules, there are about 116,000 ADP molecules in an *E. coli* cell. Alternatively, assuming an ATP concentration of 5.4 nM implies that the ADP concentration is 0.68 mM.
- MinD diffusion coefficient. MinD has a molecular weight of 29.6 kDa *(7).* Combining this with a rule-of-thumb for computing diffusion coefficients in ref. ***(8)*** gives a diffusion coefficient of 2.6 μm^2^/s. The model uses this value. It is essentially identical to the 2.5 μm^2^/s value assumed in the Huang et al. model (***9***).
- Nucleotide exchange from MinD_ADP_ to MinD_ATP_. There are two possible mechanisms for conversion from MinD_ADP_ to MinD_ATP_: transfer of just a phosphate group from free ATP to bound ADP, leading to free ADP and bound ATP, or exchange of entire nucleotides. Experiments with radiolabeled ATP showed that the latter mechanism is the correct one (***10***). There are no good *in vivo* values for the reaction rate constant for this reaction. Huang et al. (***9***) used a value of 1 s^-1^ for the reaction MinD_ADP_ → MinD_ATP_ without justification, except to note that it’s within an observed range for guanine nucleotide exchange that spans 5 orders of magnitude; note that this is a pseudo-first order reaction because it ignores the ATP reactant. However, their model behavior agrees well with the experimental data, which does provide some justification for their parameters. Lacking better values, this model assumes a rate constant of 1 s^-1^. Dividing by the ATP concentration of 1.54 mM gives a reaction rate constant of 650 M^-1^s^-1^ for the second order reaction MinD_ADP_ + ATP → MinD_ATP_ + ADP. After unit conversion, this is 1.1×10^-6^ μm^3^s^-1^. The model assumes this value for the nucleotide exchange from MinD_ADP_ to MinD_ATP_, represented here as *k_DT_* = 1.1×10^-6^ μm^3^s^-1^.
- Nucleotide exchange from MinD_ATP_ to MinD_ADP_. Lackner et al. showed that ATP competes about 3 times more effectively at binding to MinD than does ADP (their Figure 2B) (***11***). This contrasts crystal structure results, in (***12***), which suggests similar binding affinity. Based on the former result, this model assumes that the reaction rate constant for MinD_ATP_ + ADP → MinD_ADP_ + ATP is one third that of the reaction in the other direction, making it *k_TD_* = 0.37×10^-6^ μm^3^s^-1^.
- Nucleotide gain. There are no good data on the rate of nucleotide binding to unbound MinD in the reactions MinD_apo_ + ATP → MinD_ATP_ and MinD_apo_ + ADP → MinD_ADP_. Thus, this model assumes the same reaction rate constants as for the respective nucleotide exchange reactions. In particular, it assumes a rate constant of *k_AT_* = 1.1×10^-6^ μm^3^s^-1^ for the former reaction and *k_AD_* = 0.37x10^-6^ pm^3^s^-1^ for the latter reaction.
- MinD_ATP_ nucleotide loss. To my knowledge, all published models of the Min system assume that all of the cell’s MinD protein is bound to either ADP or ATP (e.g. refs. (***9,13, 14***)). This model modifies this assumption slightly, assuming instead that the equilibrium concentration of MinD_ATP_ is 20-fold higher than that of MinD_apo_. The reaction MinD_apo_ + ATP ↔ MinD_ATP_ is at equilibrium when *k_AT_*[ATP][MinD_apo_] = *k_TA_*[MinD_ATP_], using *k_AT_* and *k_TA_* as the forward and reverse reaction rate constants. From the prior numbers and the 20-fold concentration difference assumption, *k_TA_* = 0.05 s^-1^.
- MinD_ADP_ nucleotide loss. At equilibrium, nucleotide exchange between MinD_ATP_ and MinD_ADP_ is in balance, so [MinD_ATP_]/[MinD_ADP_] = *k_DT_*[ATP]/(k_TD_[ADP]). Also, ATP addition and removal is in balance, so [MinD_apo_]/[MinD_ATP_] = *k_TA_*/(*k_AT_*[ATP]). Multiplying these equations gives [MinD_apo_]/[MinD_ADP_] = *k_DT_k_TA_*/(*k_TD_k_AT_*[ADP]). Similarly, for the reaction MinD_apo_ + ADP ↔ MinD_ADP_ is at equilibrium when *k_AD_* [ADP][MinD_apo_] = *k_DA_*[MinD_ADP_], meaning that [MinD_apo_]/[MinD_ADP_] = *k_DA_*/(*k_AD_*[ADP]). Setting these two results for the same ratio equal to each other and simplifying shows that *k_DA_* = *k_DT_k_TA_k_AD_*/(*k_TD_k_AT_*). Plugging in the numbers from above, *k_AT_* = 0.05, which is the same as the *k_TA_* value.
- MinD_ATP_ dimerization and dissociation rate constants. MinD_ATP_ has been shown to be predominantly dimeric when its concentration is above 2 μm, and primarily monomeric at lower concentrations (***15***). This model assumes this value for the MinD_ATP_ dissociation constant. There are no good data for the dimerization and dissociation rate constants, so this model assumes that the dissociation rate constant is 1 s^-1^ (making it the same as nucleotide exchange from MinD_ADP_ to MinD_ATP_). Combining this assumption with the dissociation constant implies that the association rate constant is 5×10^5^ M^-1^s^-1^, which is 8.5×10^-4^ μm^3^s^-1^.
- Other dimer dissociation rate constants. This model assumes the same dissociation rate constant for all MinD dimers. From above, this is 1 s^-1^.
- MinD_ATP_ hydrolysis rate constant. The ATPase activity of MinD protein with excess ATP has been shown to be about 2.5 nmoles of ATP per mg of protein per minute ***(16).*** Converting units gives this rate constant as 1.2×10^-3^ s^-1^.

#### Smoldyn input file

~~~
# Smoldyn configuration file for MinD dimerization and nucleotide dynamics
# Units are microns and seconds

define ATPC    930000
define ADPC    116000
define KAD     (0.37e-6)*ADPC
define KDA     0.05
define KAT     (1.1e-6)*ATPC
define KTA     0.05
define KDT     (1.1e-6)*ATPC
define KTD     (0.37e-6)*ADPC
define KDIMER  8.5e-4
define KDISS   1
define KHYDRO  1.2e-3

# Graphical output
graphics opengl
graphic_iter 100

# System space and time definitions
dim 3
boundaries x 0 1 p
boundaries y 0 1 p
boundaries z 0 1 p

time_start 0
time_stop 30>
time_step 0.001

# Molecular species and their properties
species A D T

difc all 2.6
color A blue
color D green
color T red

display_size all 2

# Reactions
reaction_rule rxnAtoD *A* <-> *D* KAD KDA
reaction_rule rxnAtoT *A* <-> *T* KAT KTA
reaction_rule rxnDtoT *D* <-> *T* KDT KTD
reaction_rule rxndimer T + T -> TT KDIMER
reaction_rule rxndissoc ?? -> ? + ? KDISS
reaction_rule rxnhydro ?&T -> ?&D KHYDRO

#expand_rules on-the-fly
expand_rules all

# initial molecules
mol 2000 A u u u

output_files MinDdimerout.txt
output_format CSV
cmd B molcountheader MinDdimerout.txt
cmd i 10 30 1 molcount MinDdimerout.txt

end_file
~~~

### 5 Polymerization

#### Theory for polymer length distribution

Consider the polymerization model “polymer_end1,” in which polymerization and depolymerization arise through assembly and disassembly at one polymer end. It is expressed using the reaction rule: “A + * -> A*”. This section shows that the equilibrium length distribution is exponential.

In this model, the association and dissociation rate constants, given here as *k_f_* and *k_r_,* respectively, are the same for all polymer lengths. Based on this, define the association constant as

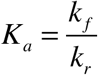

This is also the equilibrium constant for the dimerization reaction

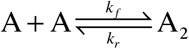

where the dimer is expressed here as A_2_, rather than as AA as it is in the simulation. As an equation, this means that

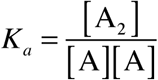

where brackets denote concentrations. The same equilibrium constant applies to longer polymers too due to the assumption that the reaction rate constants are the same for all polymer lengths. Thus,

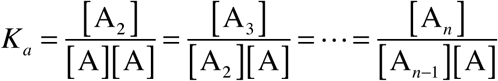

Rearrangement leads to

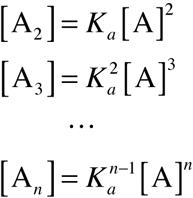

In addition to the prior assumptions, suppose the total concentration of polymer units is fixed at [A_*tot*_]. This is equal to

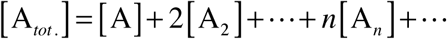

Substituting in the prior result and simplifying leads to

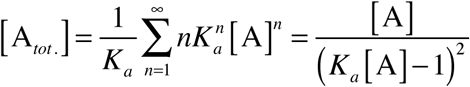

where the latter equality follows from a standard summation identity. Expanding and rearranging the latter equality leads to a quadratic equation in [A],

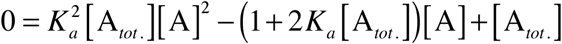

The solutions are

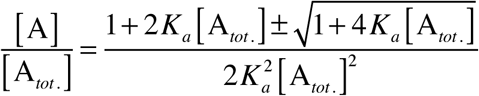

To determine whether the positive or negative root is the correct one, consider the case with low association, where *K_a_*[A] << 1. From above, this implies that [A_*n*_]/[A] << 1 for all *n* > 1 and also that [A] ≈ [A_*tot*_]. In this case, the square root in the quadratic equation can be expanded using a Taylor series, leading to

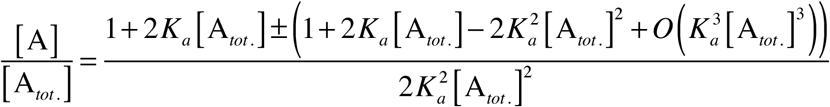

The solution with the positive sign simplifies to

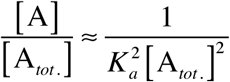

The value of the right hand side is much greater than 1 for this low association case, which disagrees with the statement made earlier that [A] ≈ [A_*tot*_], implying that this is the incorrect solution. The solution with the negative sign simplifies to

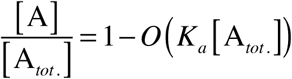

This agrees with prior statements, implying that the negative sign leads to the correct solution.

Thus, the equilibrium distribution of polymer lengths can be calculated from the two following equations, both of which are copied from above,

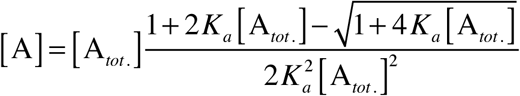

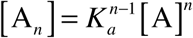

This shows that the length distribution depends exponentially on the polymer length, in agreement with prior results by Flory (***17***).

Next consider the model “polymer_end2”, which is identical to the one just shown except that monomer dimerization, through the reaction |A |+ A → A_2_, has twice the association reaction rate. Using the same definition for *K_a_* as before,

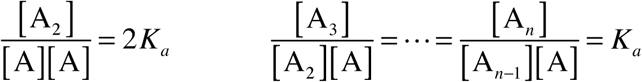

In the latter equation, *n* can adopt values of 2 or larger. Following the same procedure as before, these rearrange to

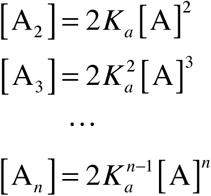

Substituting into the [A_*tot*_] definition leads to

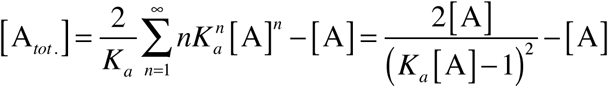

Expanding and rearranging the latter equality leads to a cubic equation in [A], which is too complex to solve here. However, importantly, the length distribution is still exponential for *n* ≥ 2,

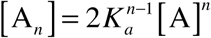

Also, the actual monomer concentration, [A], is half of the value that would be predicted from this equation.

The “polymer_mid” model is more difficult to evaluate because it has many more reactions. Of these reactions, all of the forward reaction rate constants are the same; they are all twice as large as the values in the “polymer_end1” model due to the fact that association can happen with either reactant ending up on the “left” side of the product. Also, all of the reverse reaction constants for asymmetric dissociation are the same; they are also twice as large as the values in the “polymer_end1” model, in this case because there are two ways for each asymmetric dissociation to arise. However, the reverse reaction constants for symmetric dissociation are a factor of two smaller than the others because each of these dissociations can only happen in one way. There are many fewer of these latter reactions than the others, so nearly all of the reaction equilibrium constants are the same as for the “polymer_end1” model, which also leads to the same equilibrium polymer length distribution. Figure 3 of the main text shows this result. Nevertheless, the slower dissociations for symmetric species undoubtedly affect the length distribution, and this should be particularly noticeable for small polymers because symmetric dissociations are a larger fraction of their total dissociations. Indeed, the simulation data shown in the figure shows a small but statistically significant deviation away from the exponential line for small polymer sizes, presumably arising from this effect.

The standard deviation for the equilibrium populations of different polymer lengths can be computed from the standard result that, when at equilibrium, the probability density for the population size for any individual chemical species obeys a Poisson distribution. A Poisson distribution has the variance equal to its mean, so the standard deviation is the square root of the mean.

#### Polymer length distribution figure

The following figure shows equilibrium polymer length distributions for all three models tested here. As in Figure 3 of the main text, red dots represent the “polymer_end1” model, blue dots represent the “polymer_mid” model, the solid black line represents the theoretical exponential distribution, and the dashed black lines represent the theoretical computation for one standard deviation away from the exponential distribution. In addition, the green dots represent the “polymer_end2” model. These points agree qualitatively with the predictions described above.

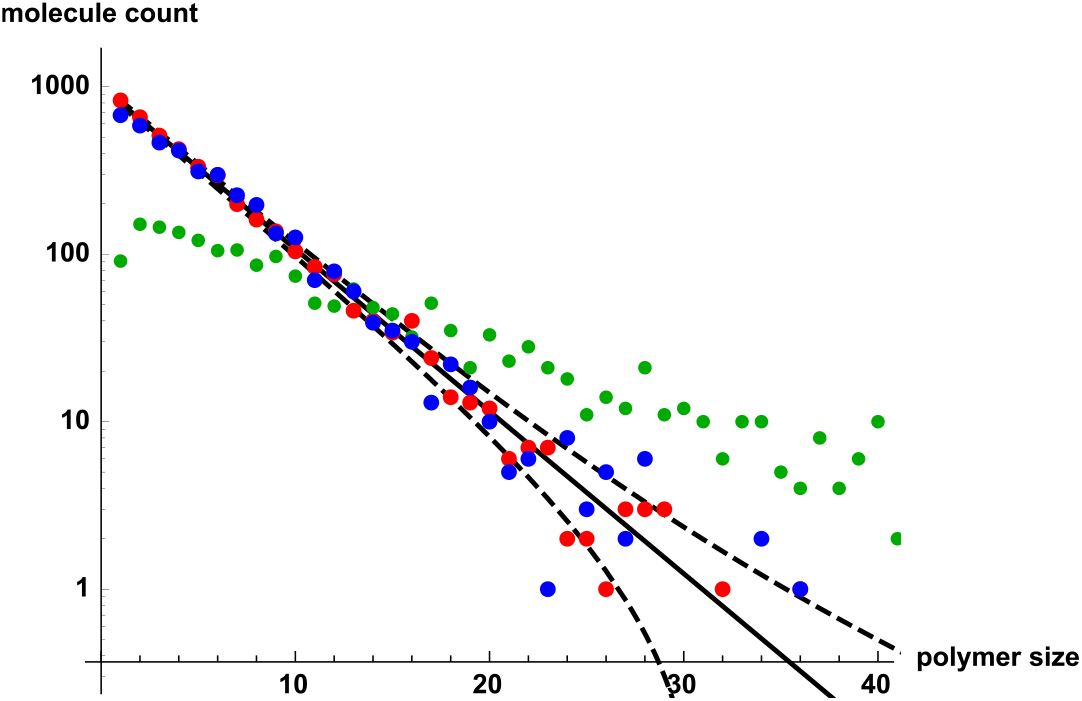

#### Smoldyn input file for polymer end1 model

~~~
# Smoldyn configuration file to test wildcards in reactions
# This file simulates polymerization, one unit at a time
# This file uses a simple reaction rule, which gives a simple length distribution
# but ignores multiplicity in the monomer association reaction.

define FWDRATE 0.1
define REVRATE 0.1

# Graphical output
graphics opengl
graphic_iter 1000

# System space and time definitions
dim 3
boundaries x 0 10 p
boundaries y 0 10 p
boundaries z 0 10 p

time_start 0
time_stop 1000
time_step 0.01

# Molecular species and their properties
species A
difc A 1
color A red
display_size A(all) 2

# Reactions
reaction_rule rxn * + A <-> *A FWDRATE REVRATE

expand_rules on-the-fly

# This reaction rule has the same reaction rate for all reactions. This may be
# correct, depending on the model. However, it is probably more accurate for the
# the monomer association reaction to be twice as fast, because its association can
# happen in either of two ways. This change, in polymer_end2.txt, makes the model
# consistent with tbe BioNetGen model of essentially the same name.

# initial molecules
mol 20000 A u u u

cmd A diagnostics all

output_files polymer_end1out.txt stdout
output_format csv
cmd N 1000 molcount polymer_end1out.txt
cmd N 1000 molcount stdout

end_file
~~~

#### Smoldyn input file for polymer end2 model

~~~
# Smoldyn configuration file to test wildcards in reactions
# This file simulates polymerization, one unit at a time
# This file uses more complex reaction rules, which is slightly less simple to analyze
# but correctly accounts for multiplicity in the monomer reaction rate.

define FWDRATE 0.1
define REVRATE 0.1

# Graphical output
graphics opengl
graphic_iter 1000

# System space and time definitions
dim 3
boundaries x 0 10 p
boundaries y 0 10 p
boundaries y 0 10 p

time_start 0
time_stop 4000
# polymers are longer here than in polymer_end1, so equilibration takes longer time_step 0.01

# Molecular species and their properties
species A
difc A 1
color A red
display_size A 2

# Reactions
reaction_rule rxn1 A + A <-> AA 2*FWDRATE REVRATE
reaction_rule rxn2 *AA + A <-> *AAA FWDRATE REVRATE

expand_rules on-the-fly

# This could also be represented with the sole rule * + A <-> *A. However, that has
# the same reaction rate for all reactions. Here, the monomer association reaction
# is twice as fast, working on the assumption that its association can happen in either
# of two ways, which makes it consistent with tbe BioNetGen model of the same name.

# initial molecules
mol 20000 A u u u

cmd A diagnostics all

output_files polymer_end2out.txt stdout
output_format csv
cmd N 1000 molcount polymer_end2out.txt
cmd N 1000 molcount stdout

end_file
~~~

#### Smoldyn input file for polymer mid model

~~~
# Smoldyn configuration file to test wildcards in reactions
# This file simulates polymerization, where any two polymers can join end-to-end,
# or any polymer can divide at any place.

define FWDRATE 0.1
define REVRATE 0.1

# Graphical output
graphics opengl
graphic_iter 1000

# System space and time definitions
dim 3
boundaries x 0 10 p
boundaries y 0 10 p
boundaries z 0 10 p
time_start 0
time_stop 1000
time_step 0.01

# Molecular species and their properties
species A
difc A 1
color A red
display_size A(all) 2

# Reactions
reaction_rule rxn * + * <-> ** 2*FWDRATE REVRATE

# the forward reaction rate is multiplied by 2 because the wildcards only consider a
# single possible bond with a reaction, whereas the reaction allows two possible bonds,
# which are on the left and right sides of the first reactant.

expand_rules on-the-fly

# initial molecules
mol 20000 A u u u

output_files polymer_midout.txt stdout
output_format csv
cmd N 1000 molcount polymer_midout.txt
cmd N 1000 molcount stdout

cmd A diagnostics all

end_file
~~~

### 6 Transcription and translation

The model here uses numbers that are approximately correct for a *Saccharomyces cerevisiae* yeast cell. In particular, it uses a spherical cell geometry with a plasma membrane radius of 2.5 μm and a nuclear radius of 1 μm; typical cell values are 2.5 to 5 μm and 0.9 μm (***18***), respectively. The simulation runs for 10,000 s, which is about 2.8 hours; typical cell generation times are around 1.5 to 2.3 hours in rich media (***19***).

The simulation assumed that diffusion coefficients were: 4 μm^2^/s for ribosomes, 0.1 μm^2^/s for DNA, 8 μm^2^/s for RNA, and 22 μm^2^/s for proteins. The ribosome value of 4 μm^2^/s is from equations and advice in Note 3 of ref. (***8***) along with the 3200 kDa molecular weight of a ribosome. The DNA value of 0.1 μm^2^/s is a very rough approximation. If the DNA sequence in the model were the complete DNA strand, then it would have a much faster diffusion coefficient (***20***) but if it is assumed that the DNA is actually a small portion of a much larger chromosome, then it would not diffuse freely. Instead, it would diffuse over a small region with so-called segmental diffusion (***21***), with a very small diffusion coefficient, likely on the order of 10^-4^ μm^2^/s (see (***1***)). Similarly, the RNA diffusion coefficient would be several-fold faster than the 8 μm^2^/s in the simulation if the simulated RNA sequences were the entire molecule and it diffused as a sphere (based on ~500 Da/basepair and equations in Note 3 of ref. (***8***)). However, yeast mRNAs typically include untranslated regions and poly(A) tails which substantially increase their molecular weights. Finally, the protein diffusion coefficient of 22 μm^2^/s is taken from the value for GFP diffusion in eukaryotic cytoplasms (***22,23***).

#### Smoldyn model

~~~
# Smoldyn configuration file to test wildcards in reactions
# Units are microns and seconds, numbers are approximately correct for a yeast cell

define KTRANSC 0.01 # 100 seconds per transcript
define KRIBBIND 0.1 # chosen to give ~100 proteins
define KTRANSL 2 # 2 amino acids per second, should be 20
define KMUT 0.000002 # 0.02 mutations per ((10000 s)*basepair*option)
define KRNADEG 0.01 # RNA lifetime of 100 seconds
define KPROTDEG 0.01 # protein lifetime of 100 seconds

define CELLRAD 2.5 # cell diameter of 5 um
define NUCRAD 1 # nuclear diameter of 5 um

random_seed 2

# Graphical output
graphics opengl
graphic_iter 1000
frame_thickness 0

# System space and time definitions
dim 3
boundaries x -CELLRAD CELLRAD
boundaries y -CELLRAD CELLRAD
boundaries z -CELLRAD CELLRAD

time_start 0
time_stop 10000 # about 2.5 hours, which is 1-2 cell generations
time_step 0.1

# Molecular species and their properties
species DnaATCAATATT Rib

difc Rib 4 # from eq. in Andrews, 2012
color Rib grey
display_size Rib 2

difc Dna* 0.1 # complete guess, assumes DNA is part of chromosome

difc_rule Dna* 0.1
difc_rule Rna* 8
difc_rule Prot* 22 # GFP diffusion coefficient

color_rule Dna* black
color_rule Rna* red
color_rule Prot* green
color_rule Rna*Rib* lightgreen

display_size_rule Dna*|Rna* 4
display_size_rule Prot* 2
display_size_rule Rna*Rib* 4

# Surfaces
start_surface cellmembrane
color both black
polygon both edge
action_rule all both reflect
panel sphere 0 0 0 CELLRAD 20 20
end_surface

start_surface nucmembrane
color both purple
polygon both edge
action_rule Dna*|Rib both reflect
action_rule Rna*|Prot* both transmit
panel sphere 0 0 0 NUCRAD 10 10
end_surface

# Compartments
start_compartment nucleus
point 0 0 0
surface nucmembrane
end_compartment

start_compartment cell
point 0 0 0
surface cellmembrane
end_compartment

start_compartment cytoplasm
compartment equal cell
compartment andnot nucleus
end_compartment

# Reactions
reaction_rule rxnTransc Dna* -> Dna$1 + Rna$1 KTRANSC

# reaction_rule rxnRibBind Rna*[A,T,C,G] + Rib -> RnaRib*[A,T,C,G]Prot KRIBBIND

reaction_rule rxnTranslN Rna*RibAA[T,C]* -> Rna*AA[T,C]Rib*N KTRANSL
reaction_rule rxnTranslF Rna*RibTT[T,C]* -> Rna*TT[T,C]Rib*F KTRANSL
reaction_rule rxnTranslL Rna*RibTT[A,G]* -> Rna*TT[A,G]Rib*L KTRANSL
reaction_rule rxnTranslS Rna*RibTC?* -> Rna*TC?Rib*S KTRANSL
reaction_rule rxnTranslY Rna*RibTA[T,C]* -> Rna*TA[T,C]Rib*Y KTRANSL
reaction_rule rxnTranslW Rna*RibTGG* -> Rna*TGGRib*W KTRANSL
reaction_rule rxnTranslL Rna*RibCT?* -> Rna*CT?Rib*L KTRANSL
reaction_rule rxnTranslP Rna*RibCC?* -> Rna*CC?Rib*P KTRANSL
reaction_rule rxnTranslH Rna*RibCA[T,C]* -> Rna*CA[T,C]Rib*H KTRANSL
reaction_rule rxnTranslQ Rna*RibCA[A,G]* -> Rna*CA[A,G]Rib*Q KTRANSL
reaction_rule rxnTranslR Rna*RibCG?* -> Rna*CG?Rib*R KTRANSL
reaction_rule rxnTranslI Rna*RibAT[T,C,A]* -> Rna*AT[T,C,A]Rib*I KTRANSL
reaction_rule rxnTranslM Rna*RibATG* -> Rna*ATGRib*M KTRANSL
reaction_rule rxnTranslT Rna*RibAC?* -> Rna*AC?Rib*T KTRANSL
reaction_rule rxnTranslN Rna*RibAA[T,C]* -> Rna*AA[T,C]Rib*N KTRANSL
reaction_rule rxnTranslK Rna*RibAA[G,A]* -> Rna*AA[G,A]Rib*K KTRANSL
reaction_rule rxnTranslS Rna*RibAG[T,C]* -> Rna*AG[T,C]Rib*S KTRANSL
reaction_rule rxnTranslR Rna*RibAG[G,A]* -> Rna*AG[G,A]Rib*R KTRANSL
reaction_rule rxnTranslV Rna*RibGT?* -> Rna*GT?Rib*V KTRANSL
reaction_rule rxnTranslA Rna*RibGC?* -> Rna*GC?Rib*A KTRANSL
reaction_rule rxnTranslD Rna*RibGA[T,C]* -> Rna*GA[T,C]Rib*D KTRANSL
reaction_rule rxnTranslE Rna*RibGA[A,G]* -> Rna*GA[A,G]Rib*E KTRANSL
reaction_rule rxnTranslG Rna*RibGG?* -> Rna*GG?Rib*G KTRANSL
reaction_rule rxnRibUnbind Rna*RibProt* -> Rna* + Rib + Prot* KTRANSL

reaction_rule rxnRnaDeg Rna*[A,T,C,G] -> 0 KRNADEG
reaction_rule rxnProtDeg Prot* -> 0 KPROTDEG

reaction_rule rxnMut Dna*?* -> Dna*{A|T|C|G}* KMUT

reaction_log stdout rxnMut all

expand_rules on-the-fly

# initial molecules
compartment_mol 1 DnaATCAATATT nucleus
compartment_mol 100 Rib cytoplasm # a cell really has ~1000 to 10000

output_file expressionout.txt stdout
output_format CSV
#cmd i 0 10000 10 molcountspecieslist expressionout.txt Rna* Prot*

cmd i 0 10000 10 molcountspecieslist expressionout.txt Rna*AAT* Rna* ProtINI ProtIYI
cmd A molcountheader stdout
cmd A molcount stdout

#cmd @ 100 diagnostics all

text_display time Dna* Rna* Prot*

end_file
~~~

